# Bayesian statistics show a lack of change in excitability following bihemispheric HD-TDCS over the primary somatosensory cortices

**DOI:** 10.1101/814780

**Authors:** Gallo Selene, Thijs J. Baaijen, Suttrup Judith, Fernandes-Henriques Carolina, Keysers Christian, Gazzola Valeria

**Affiliations:** Netherlands Institute for Neuroscience, an institute of the Royal Netherlands Academy of Art and Sciences (KNAW), 1105 BA, Amsterdam, The Netherlands; Faculty of Social and Behavioural Sciences, University of Amsterdam (UvA), 1001 NK, Amsterdam, The Netherlands

**Keywords:** HD-tDCS, bi-hemispheric HD-TDCS, SI, SEP, Bayesian statistic, reproducibility, robustness

## Abstract

**Background:** High Definition transcranial Direct Current Stimulation (HD-tDCS) is a method meant to explore the causal structure-function relationship of brain areas, developed to improve the spatial resolution of tDCS, but the validity of tDCS results is currently under intense debate

**Objective:** The goal of this study is to validate a new HD-tDCS protocol for bilateral modulation of the Somatosensory Cortex (SI). The new montage is meant to increase the focus of the stimulation while limiting the area of the scalp covered by electrodes. We aim to characterize the effect of the stimulation in terms of directionality, consistency and reproducibility.

**Methods:** We aim to leverage a 1 × 1 montage to most focally stimulate the primary somatosensory cortex (SI) and measure modulation via Somatosensory Evoked Potentials (SEP) triggered by median nerve stimulation.

**Results:** The results of Experiment1 suggest that our montage increases the amplitude of the SEP component N30. In Experiment2, we aim to replicate our finding and to assess the duration of the modulatory effect on N30 over time. Data from Experiment2 fails to replicate N30 modulation. A sequential Bayesian analysis performed on N30 data from both experiments indicates that the effect fluctuates across participants, without a clear homogenous directionality.

**Conclusion:** This study sets boundaries on the effect size that can be expected for this montage and illustrates the need to include replication samples or larger sample sizes to avoid overestimating effect sizes. We conclude that our montage has insufficient effect size for use in moderately sample-sized experimental studies and clinical applications.

## INTRODUCTION

The studies of Priori and his colleagues (1) followed by Nitsche and Paulus (2) have suggested that low direct electrical currents applied over the scalp can influence brain excitability and produce substantial aftereffects on cortical excitability, lasting from minutes to hours. Combined with its low cost, simple application and portability, this has led to an increasing use transcranial direct current stimulation (tDCS) in a wide variety of settings (DaSilva et al., 2015; Bikson et al., 2017b) (4–9). Unfortunately, in its standard form, using large electrodes (most commonly between 25-35 cm^2^), the effects of tDCS are too diffuse to attribute effects to specific brain regions(10,11),(12). High Definition transcranial Current Stimulation (HD-tDCS) was created to increase spatial selectivity and probe the function of specific brain regions(11,13–15). HD-tDCS uses small ring electrodes 3-5mm in diameter, with the most popular montage being a *4 × 1 ring* with a central electrode (anode or cathode) over the targeted area, surrounded (at 3–7.5 cm radius) by four reference electrodes (11,14,16–20). This increases focality (21) up to 80% compared to standard tDCS (11,20,4,20,22).

Simulations using the Finite Element Method (FEM) suggested that an increment of focality can be reached by lessening the number of return electrodes: Using only one cathode and one anode should support the most focal stimulation possible, albeit at the cost of depth current penetration (20).

Here we aim to leverage a 1×1 montage to most focally stimulate the primary somatosensory cortex (SI) and measure such modulation via Somatosensory Evoked Potentials (SEP) triggered by median nerve stimulation. Different SEP components are thought to reflect activity from different sub-regions of SI: N20 and P24 the first cortical components, from Brodmann area 3b(23–25) (25–27), N30 from precentral (28) or postcentral cortical regions (27), and P45 from area 1 or 2 (Allison et al. 1992). To modulate the hand knob of SI bilaterally, we place two anodes over the hand representation of SI (between C3 and CP3, and between C4 and CP4), and two cathodes between CZ and CP1 and CZ and CP2 (Figure 1A), so that the current flow would be parallel to the central sulcus (29). We aim to characterize the effect of the stimulation in term of directionality (increase vs. decrease of SEP), consistency (across participant) and reproducibility (across studies).

**Figure 1.**
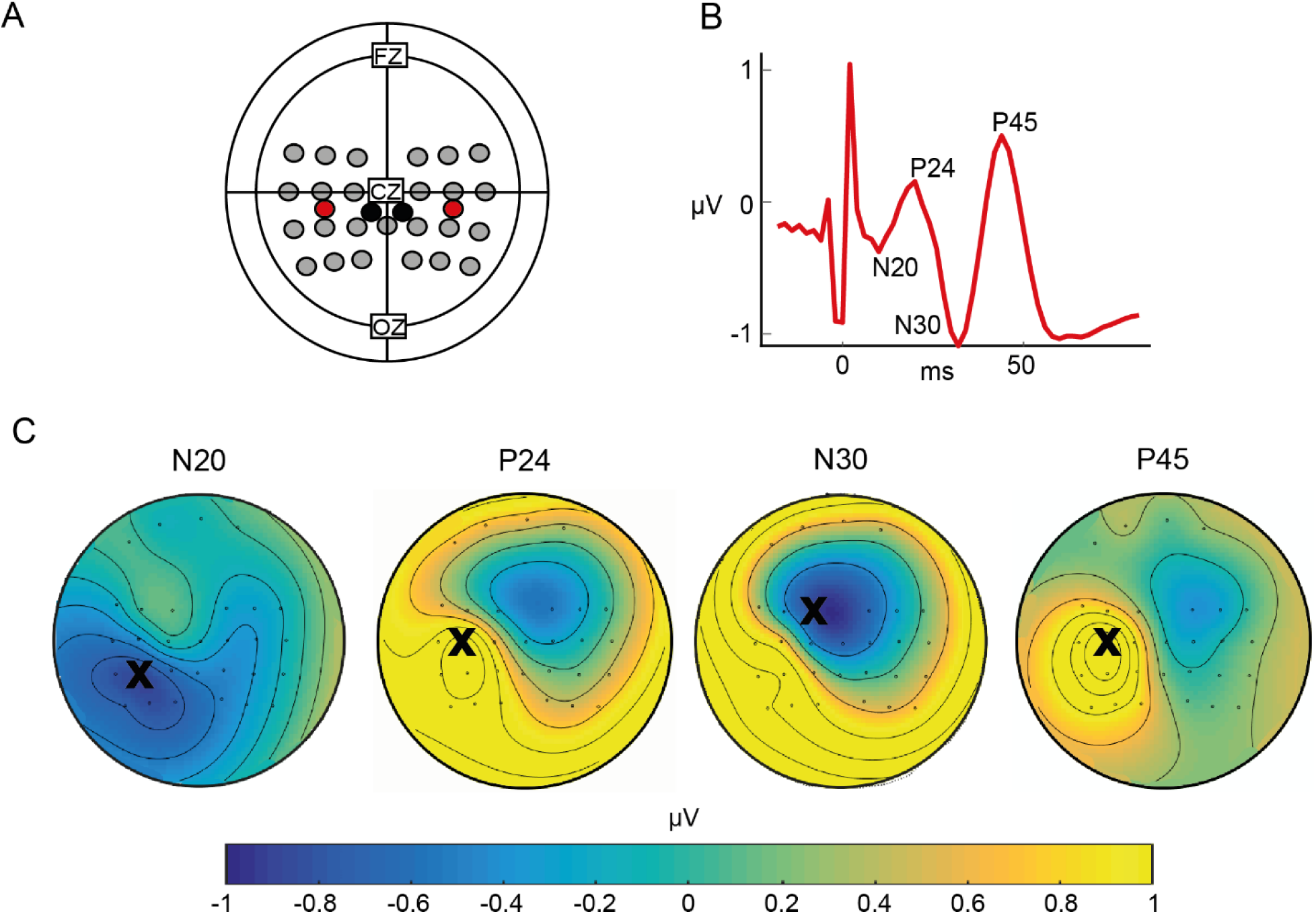
A) HD-tDCS montage and EEG electrodes. Grey circles are the EEG electrodes. Red circles represent HD-tDCS anodes, and black circles HD-tDCS cathodes. B) SEP obtained by averaging across experiments the signal recorded in C3 at T_0_, before HD-tDCS stimulation. C) Scalp topography of the SEP components obtained by averaging the T0-grand average of experiment 1 and 2 in the appropriate time-windows. Bold Xs indicate the location of the electrodes chosen as peak of the component.

In Experiment 1, we measured the change in cortical excitability in SI by comparing SEPs before and after the tDCS stimulation. A meta-analysis suggests that for tDCS over sensorimotor cortices the anode causes an enhancement of cortical excitability(30) and thus we expect an *enhancement* of the SEP components (31–33) from our montage. Because the effectiveness of tDCS is debated (34–37), we performed a second experiment (Experiment 2) aimed at replicating the results of Experiment 1, and at investigating the duration of tDCS effects.

## METHODS

### Participants

Thirty-two participants took part in Experiment 1 (17 females, average age=25±5 years old) and 17 in Experiment 2 (10 females, average age=22±3). None reported neurological, psychiatric, or other medical problems or any contraindication to brain stimulation (38,39). All were right-handed, naïve to the purpose of the experiment and free to withdraw at any point. No discomfort or adverse HD-tDCS effects were reported by participants or noticed by the experimenters. All studies have been approved by the Ethics Committee of the University of Amsterdam, the Netherlands (protocol number 2015-BC-4070 and 2014-EXT-3829).

### General protocol

Both experiments consisted of a sham and an active HD-tDCS sessions over SI distributed over two days. Each session started with the calibration of the SEP system, and the recording of a first baseline SEP measurement (t0). Participants then received sham or active HD-tDCS, depending of the pre-assigned order of the conditions, which was followed by a second SEP recording in Experiment 1 (t1), and a total of seven additional recordings in Experiment 2. To reduce the variability in participants’ state(40–42), during the HD-tDCS stimulation we showed emotionally-neutral videos and induced tactile sensations on the dorso of both hands (obtained through electrical stimulation (square wave pulses, frequency of 1 Hz, duration of 2000 µs). For the entire duration of the experiment, participants sat comfortably in an armchair with their hands resting on a pillow, in a semi-darkened room in front of the computer screen. Before leaving the participants completed a tDCS side-effect questionnaire.

### HD-tDCS stimulation protocol

HD-tDCS was applied with a 9-channel stimulator (tDCS MXN-9 HD Stimulator, Soterix Medical Inc, US). To bilaterally stimulate SI we used two high-definition 5.5 mm electrodes(43) over the right and two over the left hand knob of the somato-motor cortex (Figure 1A). One anode was placed between P3 and CP3 (left side), and one between P4 and CP4 (right). The two cathodes were placed along the central sulcus between CZ and CP1 (left), and CP2 (right). The electrode holders (HD-M, Soterix Medical Inc, US) were embedded in standard EEG caps (ActiCAP, Brain Products GmbH, Germany). In the active stimulation, the current was delivered with a ramp-up time of 30s, held at 0.75 mA in both anodes (1.5 mA in total(3)) for 18 min, and ramped down over 30s. In the sham condition, the current was ramped up and back down at the beginning and at the end of the 18 min and kept around 0 mA during this period(19). The participant’s skin was cleaned by means of a Q-tip soaked in alcohol, and gel was used as interface between the skin and the electrodes. The impedance measured before the stimulation was below 10 kΩ(14,19).

### SEP stimulation protocol

SEP stimulations were obtained via non-painful electrical stimulation (Stimulator Model DS7A, Digitimer Ltd, UK) of the median nerve at the right wrist (square wave pulses, personalized stimulus intensity, frequency of 3 Hz and duration of 500 µs). The intensity, determined beforehand in each session, was set to elicit a consistent and visible twitch of the thumb, and was kept constant before and after HD-tDCS.

An opaque panel covered participant’s hands during medial nerve stimulation, preventing a modulation of SEP response through observation (44). In Experiment 1, the SEP recording consisted of one block of 500 medial nerve stimulations right before and one after the HD-tDCS stimulation. In Experiment 2, the number of medial nerve stimulation was increased to 1000 per block, in order to better differentiate the signal from noise(45), and 8 blocks were recorded: t0 before the tDCS stimulation, t1 directly after stimulation, t2 after 15 min from the end of stimulation, t3 after 30min, t4 after 45 min, t5 after 60 min, t6 after 90 min and t7 after 120min. During the medial nerve stimulation, participants were instructed to fixate their eyes on a white cross on a black screen.

### EEG Data Acquisition and Preprocessing

Electrophysiological recordings were obtained from 27 Ag/AgCl scalp electrodes (10-20 International System; Figure 1A). The ground electrode was positioned in FPz. The impedance was kept below 5 kΩ. The signal was acquired at a 500 Hz sample rate and stored on a disk for off-line pre-processing (actiCHamp and PyCoder v1.0.8, BrainVision LLC, USA). Digitalized signal has been analysed using the FieldTrip Toolbox(46) and customized MATLAB (Mathworks Inc., Natick, MA, USA) scripts. For each SEP recording, data before the first medial nerve modulation and after the last were removed, leaving a long segment of data containing only medial nerve stimulations. Signal was demeaned, detrended and filterer between 2 and 200 Hz. Using Independent Component Analysis (ICA), eye blinks, eye movements and stimulator artefacts were identified and rejected (24/26 components kept on average). The data was then segmented in trials containing one single event, 100 ms prior to each medial nerve stimulation and 300 ms after. Trials containing muscle artefacts were identified (cut off: value 6 z point above average) and excluded from further analysis (496/500 trials kept on average) using the Fieldtrip automatic rejection routine.

ERPs of each repetition were obtained by averaging the data and correcting for baseline (average time between 50 and 30 ms before medial nerve stimulation).

## DATA ANALYSES

At the group level, the signal recorded in sham and active sessions at t0 were averaged to identify SEP components (T0-Grand Average). The early SEP components known in literature as N20, P24, N30 and P45 (Allison et al. 1991, 1992; Valeriani et al 2001) were clearly identifiable in the T0-Grand Average of both experiments. Visual inspection of the scalp topographies (Figure 1C) and the information available in literature (24,25,47) guided the selection of peak electrodes and of component time-windows. We defined N20 as the most negative peak between 18 and 22 ms after medial nerve stimulation recorded in the electrode P5, component P24 as the most positive peak between 22 and 26 ms recorded in C3, N30 as the most negative peak between 28 and 36 recorded in FC1 and P45 as most positive peak between 42 and 48 ms recorded in CP3. For illustrative purposes we averaged the T0-Grand Average (in C3) obtained from the two experiments in Figure 1B.

At the individual level, for each participant and block (time-repetition), we calculated the SEP components of interest in the electrode selected in T0-GrandAverage as the maximum or minimum peak (according to the component polarity) present in the time-windows selected at the group level (47).

### HD-tDCS effect analyses

To assess the effect of the experimental manipulation, for sham and active stimulation sessions, we subtracted, for each participant and ERP component separately, the baseline measurement (t0) from each measurement after active or sham HD-tDCS. Given in Experiment 1 we only have one SEP recording after HD-tDCS the index of stimulation effect for a component becomes: (t1_active_-t0 _active_) - (t1_sham_ -t0 _sham_). These values were used at the group level to test whether stimulation had a consistent effect across participants. Because the anodes were placed above the source of the SEP components, we expected HD-tDCS to enhance SEP components - i.e. to increase negativity for N20 and N30 and positivity for P20 and P45. As the distribution of indices did not significantly deviate from normality (Table 1, Figure 1), in Experiment 1 we tested our hypotheses with four frequentist one-tail t-tests, one for each component (H_1_ for P24 and P44: index>0; H_1_ for N20 and N30: index<0). As in many cases, this lead to non-significant t-values, to quantify evidence for null hypothesis, we complement the frequentist approach with Bayesian t-tests in Jasp 9.2 (https://jasp-stats.org/) and default Cauchy prior (r=0.707) unless otherwise specified. We first looked for evidence for the null for the one-tailed effects we expected. If the Bayesian t-test brought evidence for H_0_ (i.e. index >=0 for N20 and N30, or index <=0 for P24 and P44) a second Bayesian two-tailed t-test was performed to quantify evidence specifically for H_0_: index=0.

**Table 1.**
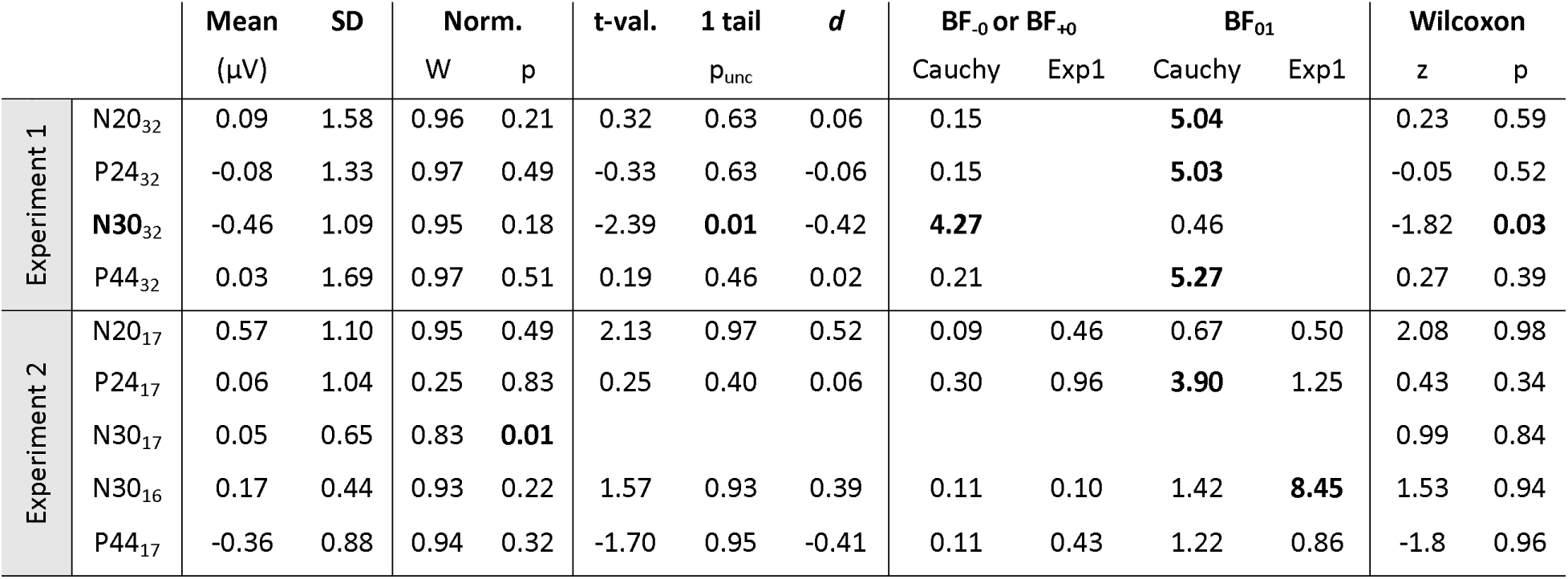
For each of the ERP component of interest the table reports: the mean value in µV of the HD-tDCS effect index [(t1active-t0 active) - (t1sham -t0 sham)]; its standard deviation (SD); the Shapiro-Wilk W- and p-values of the Normality test (Norm.); the t- and uncorrected p-value from the frequentist statistic (H1: index<0 for N20 and N30, and index>0 for P24 and P44) and the Cohen’s d effect size. Thereafter we provide the results of the one- and two-tailed Bayesian analysis. This includes the Bayes factor (BF) in favour of the expected directional effect (H_1_ index<0 for N20 and N30, and index>0 for P24 and P44, indicated with BF_-0_ and BF_+0_ respectively). We then present the Bayes factor from a two-tailed Bayesian t-test in favour of the absence of any effect (BF_01_). In all cases the Bayesian analyses has been run using the default Cauchy prior (Cauchy). For Experiment 2, BF were additionally calculated using the posterior distribution from Experiment 1 as an informed prior (Exp1). The last two columns report the z- and p-value from the non-parametric Wilcoxon signed-rank directional test (index<0 for N20 and N30, and index>0 for P24 and P44). Sample sizes indicated as subscript of each component. In bold are the significant results, and the evidence in favour of the predicted or unpredicted effects.

Using the JASP formalism, the indices next to Bayes Factors (BF) indicate what hypothesis is in the nominator and denominator. For two tailed tests, BF_10_ is p(data|H_1_)/p(data|H_0_) and BF_01_= p(data|H_0_)/p(data|H_1_). BF_10_>3 thus is moderate evidence for an effect, BF_10_<1/3 for the absence of an effect, and the reverse is true for BF_01_. BF values around 1 indicate the data is similarly likely under H_0_ and H_1_, and cannot adjudicate in favour of either. For one-tailed tests, the index representing H_1_ is indicated by a “–” if H_1_: index<0, and a “+” if H_1_: index>0. So BF_-0_ represents p(data|H_1_:index<0)/p(data|H_0_:index>=0).

Experiment 2 was collected to replicate Experiment 1 and to investigate the duration of HD-tDCS effects. The Shapiro-Wilk test for normality rejected the notion that the N30 component was drawn from a normal distribution (Table 1), which became normal after removal of participants #8. There was no significant deviation from normality for the other components. To investigate replicability of the results of Experiment 1, we first computed both a frequentist and Bayesian analysis on the index (t1_active_-t0 _active_) - (t1_sham_ -t0 _sham_) removing participant #8 for the N30 component and using all 17 participants for the other components, and then ran a non-parametric Wilcoxon signed-rank test on the full sample for each component separately. Based on Experiment 1, we expected to find an increase of negativity only for N30. To investigate whether the effect of HD-tDCS on N30 lasts over time we performed a repeated measures Bayesian ANOVA with 7 indices, one for each time point acquired after stimulation: (t*i*_active_-t0_active_) - (t*i*_sham_ -t0 _sham_) with *i*=1 to 7. We also performed Bayesian t-tests on each of these time points to assess whether they deviated from zero in the expected direction. Because the increase of negativity for N30 was not replicated in Experiment 2 for any time point, follow-up Bayesian analyses were implemented to understand the reasons for the non-replication. A sequential analysis was performed on all the four ERP components to see how the evidence in support for or against our effect accumulate over participants across the two experiments. To explore whether effects might be detectable for any of the SEP components, we also performed a 4 components x 7 time points repeated measures Bayesian ANOVA. All Bayesian ANOVAs were conducted in JASP with default priors.

## RESULTS

### Experiment 1

Traditional frequentist t-tests showed a significant increase in the negativity for N30 after HD-tDCS. No significant effects were observed for the other components (all p_unc_>0.3). A one-tailed Bayesian t-test confirms the frequentist results showing moderate evidence for an increase of negativity for N30, and indicates a moderate to strong evidence for a lack of an increase of negativity for N20, and of positivity for P24 and P44. Follow-up Bayesian tests confirm a moderate evidence for the absence of any changes in the N20, P24 and P44 components. Table 1 and Figure 2 summarize these results.

**Figure 2.**
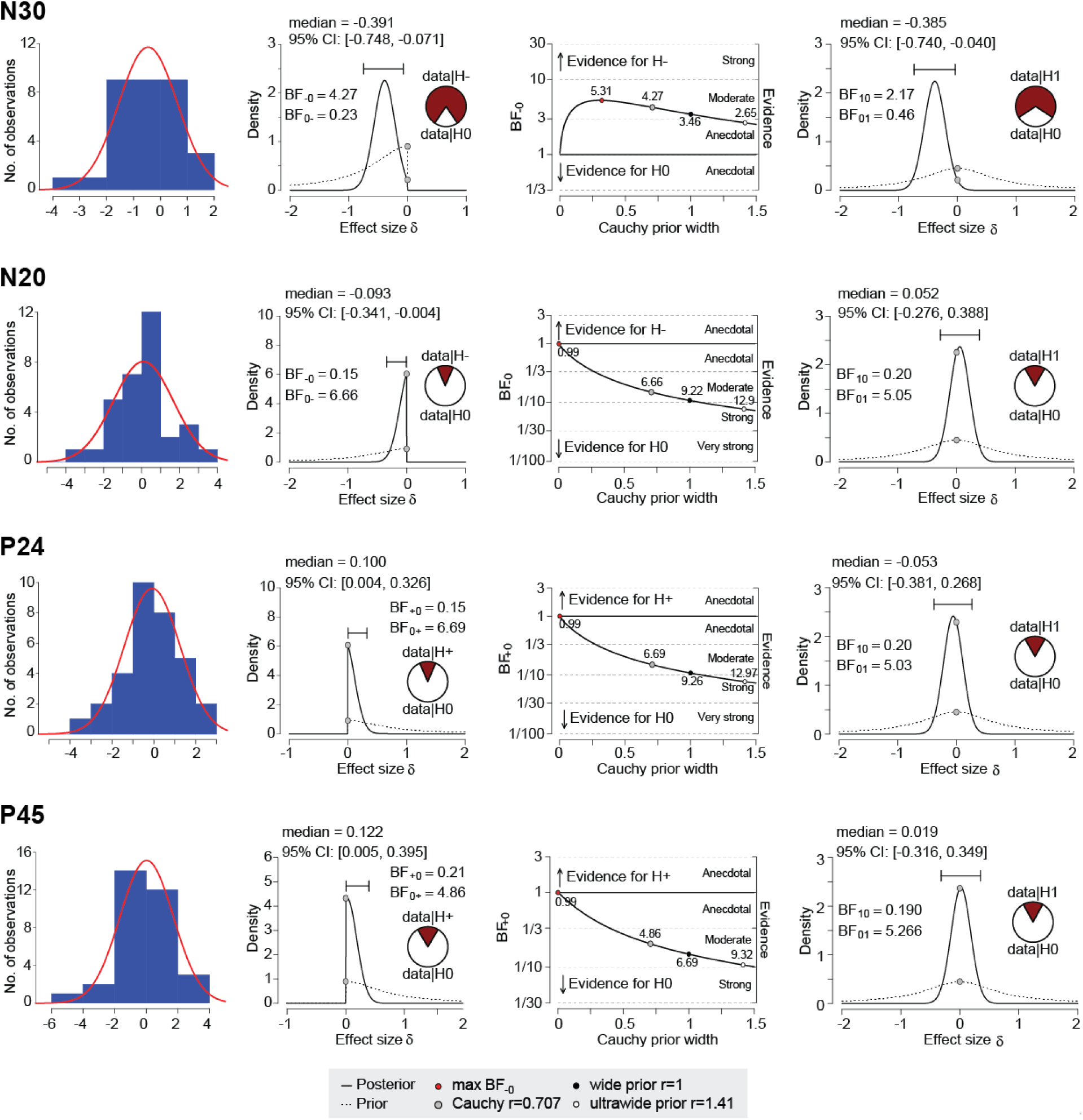
Bayesian results for Experiment 1. First column: distribution of the ERP components across participants. Second column: shape of the prior distribution (dotted line) based on our hypothesis, and the posterior resulting distribution (thick line). For Experiment 1 the default Cauchy prior was used, and the fact that the prior is set to zero on one side of each graph expresses the directionality of the hypothesis. The cake illustrates the relationship of the probability of the data given H_1_ (either an increase of negativity H_-_ or an increase of positivity H_+_) in dark red, and the probability of the data given H_0_ (i.e. either evidence for no effect or for the effect in the opposite direction). Posterior median and 95% confidence intervals (CI) of the effect size are also indicated. Third column: robustness of the Bayes Factor against modifications in the width of the prior. Fourth column: results of the two-tails Bayesian t-test. Conventions as in second column.

### Experiment 2

A frequentist one-tailed t-test on N30_16_, for which based on the results of Experiment 1 we expected to find an increase of its negativity, did not reveal significant results (Table 1). A Bayesian t-test with the default prior indicates evidence against an increase in N30 negativity (Table 1 and Figure 2). From the results of the Bayesian two-tailed test of Experiment 1 we observed that the posterior distribution was approximately normal with median at -0.39 (Figure 1, fourth column), and a SD of about 0.17 (CI/4). To take full advantage of the Bayesian framework, which allows to use of informed prior, we repeated the t-test by using a normal distribution with mean -0.39 and standard deviation 0.17 as prior. Although this prior shifts the posterior towards the expected negative effect size, results still confirm a moderate evidence for the changes in N30 negativity to be equal to zero. The non-parametric Wilcoxon signed-rank test on the full N30_17_ data set from Experiment 2 confirms the lack of reproducibility of a significant decrease in negativity for N30 (Table 1).

For the other components, frequentist and non-parametric analyses show a consistent lack of significant difference between tDCS and sham stimulation, and Bayesian results consistently show a lack of evidence in support of a change after tDCS stimulation (Table 1).

Considering our initial aim to test whether the increase of negativity in Experiment 1 would last over time, despite not replicating the effect of HD-tDCS on N30 at t1 in Experiment 2, we took advantage of the multiple recordings to test whether the reduction in negativity would be present at any other time points. A Bayesian repeated measure ANOVA on N30_16_ indicates evidence for a lack of difference between time points (the variable time as a factor has a BF_inclusion_=0.23, providing moderate evidence against inclusion of time). Results of Bayesian t-tests ran for each time point separately finally indicate that there is no evidence for an increase in N30 negativity following tDCS stimulation at any time points recorded during Experiment 2 (Figure 4 and Table 2). The non-parametric Wilcoxon signed-rank test performed on the full sample separately for each time point supports the Bayesian results, indicating that none of the time point significantly differs from zero (all p>0.16).

**Table 2.**
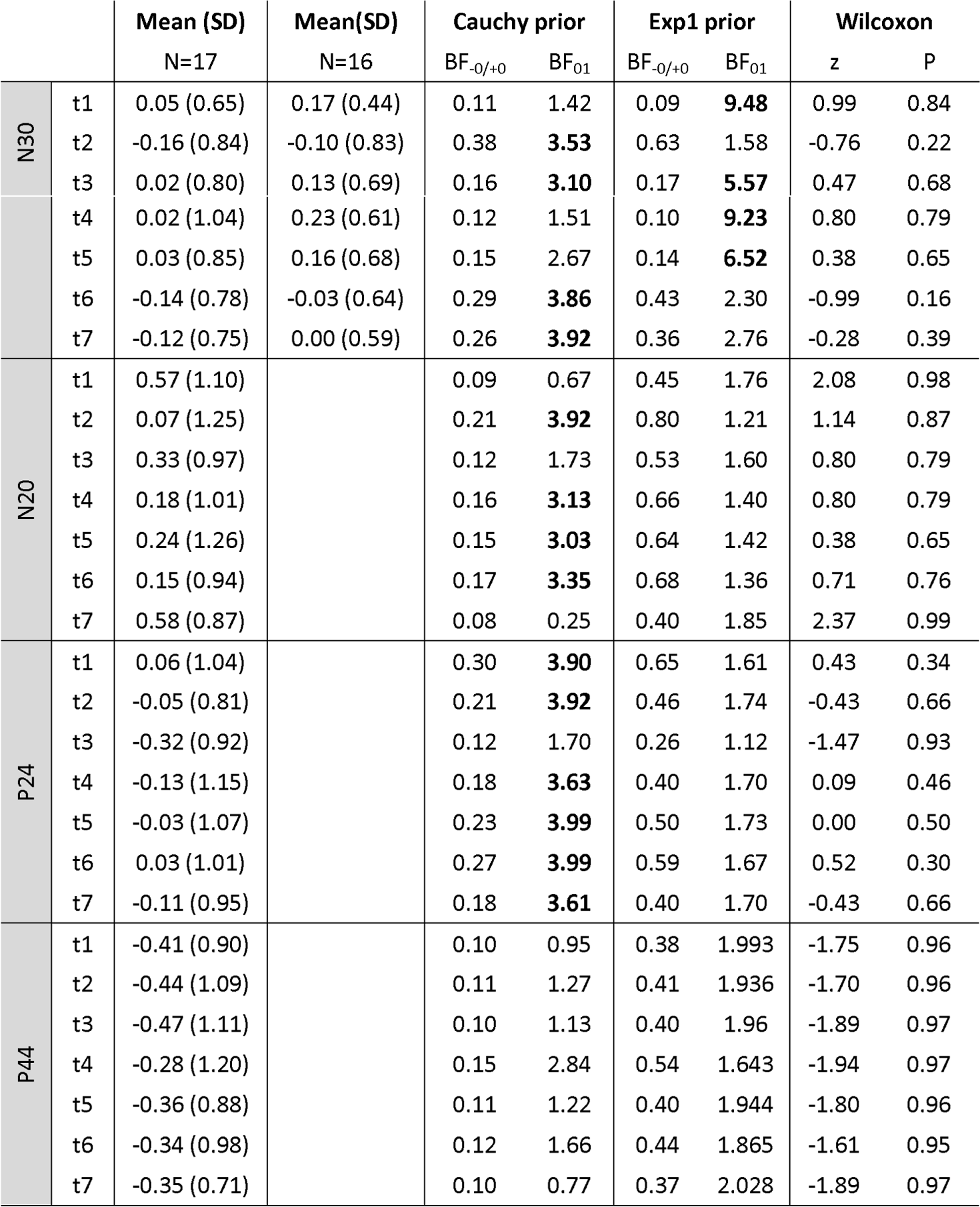
Results of Experiment 2 for each time point. For each ERP component and time point the table reports: the mean and standard deviation (SD) of the HD-tDCS effect index; the Bayesian factors calculated either using the default Cauchy prior or the prior informed by the posterior distribution calculated in Experiment 1 for t1.(BF_-0/+0_ indicates the evidence for an increase of negativity of N30 or N20 and of increase in positivity for P24 and P44; BF_01_ indicates the evidence for a null effect); z- and p-values of the non-parametric tests on the full sample. For N30 the mean and SD are calculated for the full, N=17, and partial, N=16, sample; the BF on N=16 and the non-parametric test on N=17. For all other components N=17.

**Figure 3.**
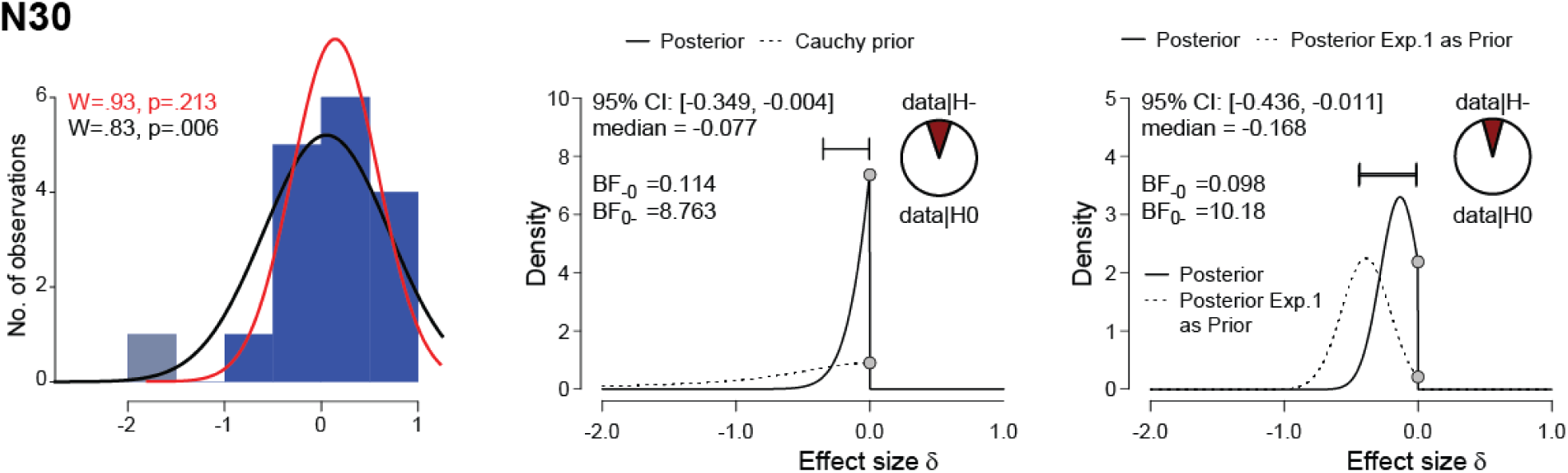
Bayesian results for Experiment 2. Distribution of the N30 components values in Experiment 2 with the normality curve fitted for the whole sample (N=17) in black and after removal of participant 8 (grey bar) in red (left graph). The middle and right most graphs show the posterior and prior distribution for the N30 component computed removing participant 8 using a default Cauchy distribution as prior (middle graph) or the posterior distribution resulting from the data of Experiment 1 (right graph). Conventions as in Figure 2.

**Figure 4.**
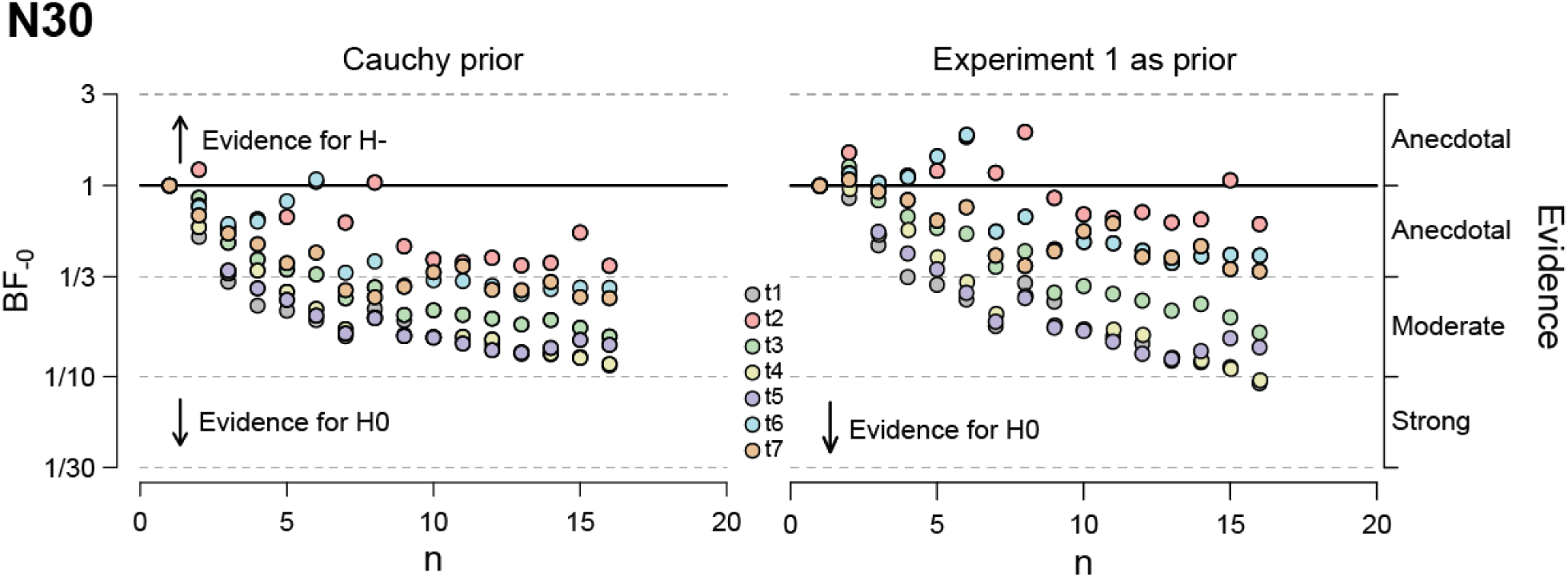
Bayesian sequential analysis for different time points. The graphs illustrated cumulative updates in BF estimation as individual participants are added to the analysis for the N30 ERP component in Experiment 2. The computation is done separately for each time point after stimulation (t1 to t7). Left: estimation calculated using the default Cauchy prior. Right estimation computed using the posterior distribution from Experiment 1 at t1 as informed prior.

To test if there is any evidence that tDCS transiently altered any of the SEP components, we performed a Bayesian repeated measure ANOVA including all the time points for each ERP component (Component x Time Point as within subject factors). This revealed that the best performing model was the one only including the factor component (BF_M_=1269). The BF for including the factor time or the component x time interaction are BF_inclusion_=0.002 and BF_inclusion_=0.000015 showing that the data provides strong evidence against the hypothesis that indices calculated at subsequent time points different from each other. The results of follow up Bayesian t-test and Wilcoxon signed rank test (Table 2) all confirm the lack of a tDCS effect at any time point.

To check whether the lack of reproducibility of the effect for the N30 component was due to a general difference in the distribution of our indices between the two experiments, we ran a Bayesian ANOVA with the four components as repeated measures and experiment as between subject factor. The ANOVA indicates that the best model is the null model (BF_M_=6.12), with BF in favour of the inclusion of the factors all being less than 0.21. A non-parametric comparison of the N30 indices across experiments also indicates a lack of significant difference across experiments (p=0.1, W=-1.63).

To have a better understanding of our data, and of why the effect found in Experiment 1 did not replicate, we pooled the data of the two experiments together and ran a sequential analysis, which calculates the evidence for H_1_ sequentially (and cumulatively) for each added participant: results at n=x consider the data including the first x participants, results at n=x+1 those including the first x+1 participants, and so forth. Figure 6 shows that while for N20, P24 and P44 the evidence in favour of a null effect keeps accumulating over participants in a relatively constant manner reaching a moderate (for P24 and P44) to strong (for N20) strength, this is not the case for N30. The sequential analysis on N30 indicates that the effect fluctuates across participants, without a clear homogenous directionality. More specifically while some participants contribute toward the predicted effect (increase in the N30 negativity), others contribute to the evidence in the opposite direction, leaving the evidence in favour of the increase of negativity anecdotal at best. In particular, this analysis shows that for N30, we happened to stop collecting data in Experiment 1 at the peak of the evidence. To be noted that the distribution of the N30 components starts to violate normality after participants 37 (32 subjects from Experiment 1 + the first 5 participants of Experiment 2; from participant 38 the Shapiro-Wilk test for normality W=.94 with p=.042), making the interpretation of this analyses for N30 more tentative. Table 3 reports the results of a 2-tailed Bayesian t-test, which are useful for planning future experiments.

**Table 3.** Effect size estimates from full sample. The table indicates for each component the median effect size (Cohen d) of the index at t1 together with the 95% confidence interval (CI) as resulting from a Bayesian t-test, together with the Bayes factors of a Bayesian 2-sided t-test (indices different from 0). Considering that the distribution of the N30 indices becomes not normal from participant 38, the values are reported both for the full not-normal sample (N30_49_) and the reduced normal sample (N30_37_). Note that in all cases effect sizes are small at best, and CI always include the zero line, suggesting that there is no confidence that the effect size differs from zero.

| | Median | 95% CI | $BF_{10}$ | $BF_{01}$ |
| --- | --- | --- | --- | --- |
| **N20** | 0.17 | -0.10, 0.44 | 0.32 | 3.13 |
| **N30<sub>49</sub>** | -0.27 | -0.55, 0.01 | 1 | 1 |
| **N30<sub>37</sub>** | -0.32 | -0.65, 0.01 | 1.24 | 0.81 |
| **P24** | -0.02 | -0.3, 0.24 | 0.16 | 6.36 |
| **P45** | -0.07 | -0.34, 0.21 | 0.18 | 5.72 |

**Figure 5.**
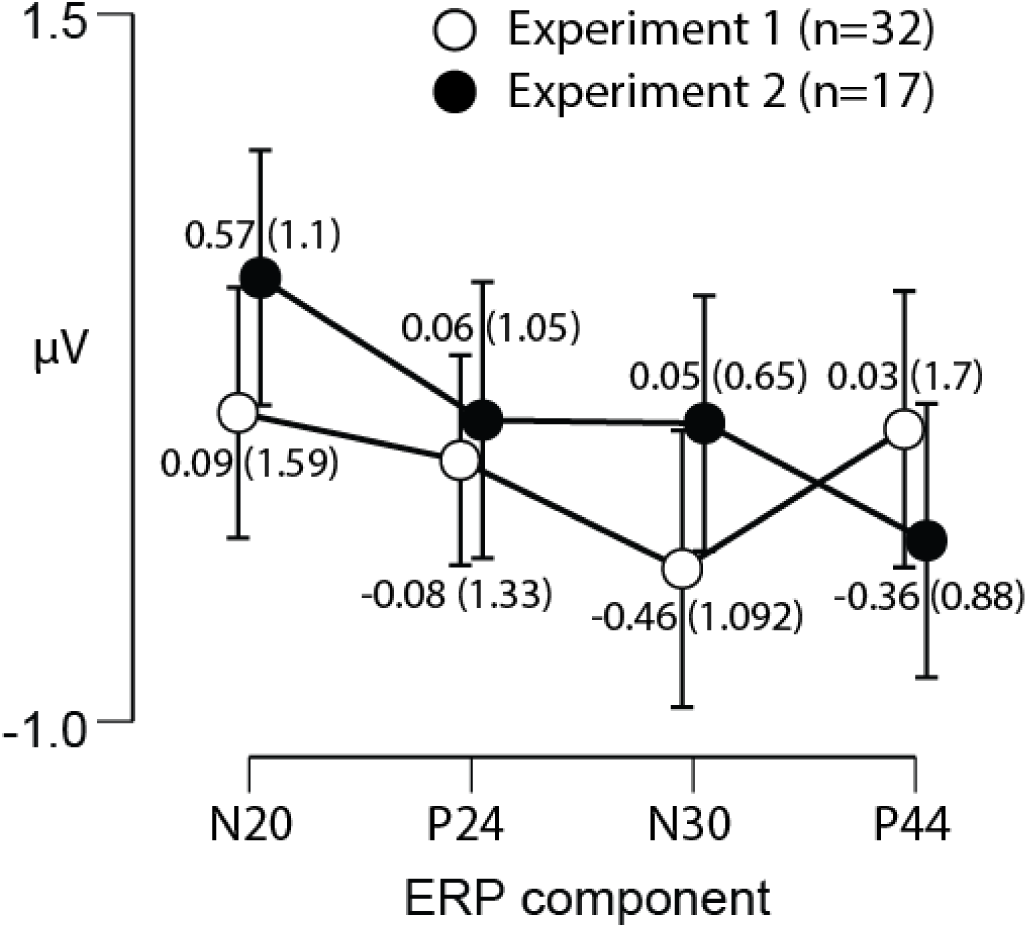
Experiment 1 and 2 comparison. The graph reports the mean indices [(ERPcomponent_t1_ active-ERPcomponent_t0_ active) - (ERPcomponent_t1_ sham - ERPcomponent_t0_ sham)] for each ERP component and the two Experiment separately. The numbers close to the dots indicate the mean and standard deviation values, while the error bars are the 95% confidence interval.

**Figure 6.**
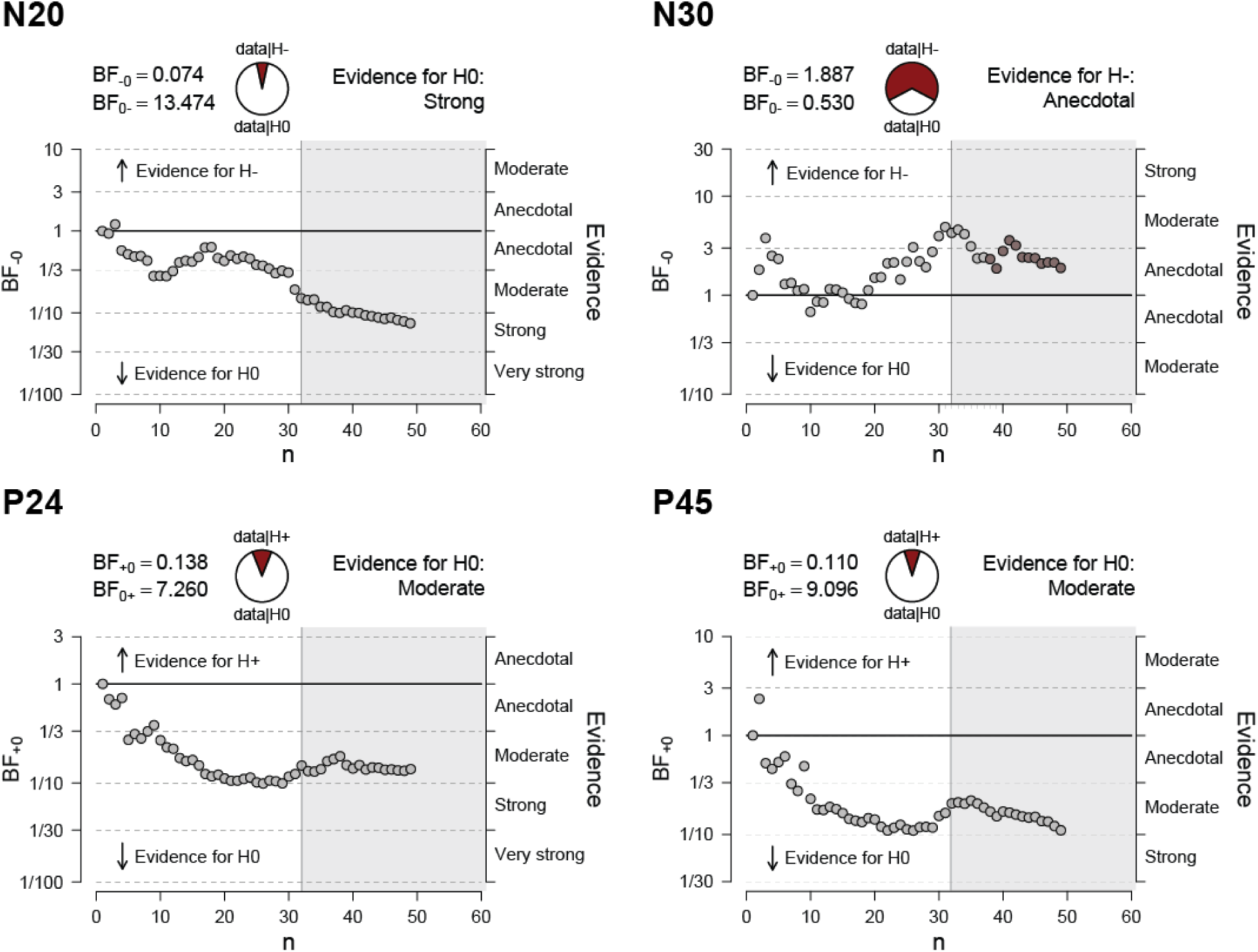
Bayesian sequential analyses. Cumulative probabilities of the data given our hypotheses for each added participant. The grey area indicates the pool of participant coming from Experiment 2. The beginning of darker dots in the N30 plot indicates that from that point onward (n=38) the N30 distribution violates normality (Shapiro-Wilk W=.94 with p=.042).

## DISCUSSION

After initial enthusiasm, the effectiveness of currently used tDCS protocols is increasingly debated(34,37,48–51,52,53,54,55). Recently, a protocol of good practices for rigor and reproducibility in tDCS research has been published(35). We designed our studies to conform to those guidelines and tried to replicate our own results. Even though Experiment 1 results seemed to suggest that our montage modulated S1 excitability, the effects of the stimulation were small (Cohen d estimated around 0.4) and characterized by individual variability. A lack of reproduction of the effect in Experiment 2 would perhaps not be entirely alarming, as the smaller study would have been expected to detect an effect size of 0.4 in only half the cases (power analysis, n=17, d=0.39, alpha=0.05, power=0.46). Bayesian sequential analyses though revealed that the results at the end of Experiment 1 were a singular peak of significance in a set of data. This points towards such a small effect size (d=0.27) that one would need to conduct studies with 87 participants to achieve 80% at alpha=0.05. As a result, this kind of tDCS protocol would be highly impractical in most cognitive neuroscience or clinical contexts.

The bias towards publishing positive results in the scientific literature can lead to a substantial literature with limited evidential value(53). This problem is particularly felt in the non-invasive brain stimulation field. The widespread use of frequentist statistic means one can provide evidence only for the presence of an effect, but not its absence. While p<0.05 arguably provide publishable evidence for a role of a brain region in a task, non-significant P-value do not translate into a likelihood that the null hypothesis is true(56,57), i.e. that a region is not involved in a task. A series of articles have stressed these limitations and proposed alternatives(58–63). The Bayes factor hypothesis testing in particular allows researchers to quantify the relative evidence for and against the null hypothesis, and to monitor it continually as data accumulate(64,65). In this study, after identifing a HD-tDCS modulation of N30 in Experiment 1 through frequentist approach, one could have been tempted to publish the result, leading others to invest in HD-tDCS to study the increasingly recognized function of SI in higher cognitive functions (66–68). Since we did not replicate the results in Experiment 2, we took advantage of the Bayesian framework. The sequential analysis on N30 indicates that the effect fluctuates across participants, without a clear homogenous directionality, with the contribution of some participants moving the evidence toward the predicted effect (increase in the N30 negativity), and others’ contributing to the evidence in the opposite direction, leaving the evidence in favour of the increase of negativity at best anecdotal, and effect sizes clearly too small to be of much use in cognitive neuroscience. In particular, this analysis shows that for N30, we happened to stop collecting data in Experiment 1 at the height of the evidence. Our montage thus produces insufficient effect size for use in moderately sample-sized experimental studies and clinical applications.

We designed the HD-tDCS montage tested in this study with the goal of maximizing the focality of the stimulation over SI. The risk of increasing focality is to loose current penetration(4,20,22). Intuitively, this is because reducing the distance between anode and cathode reduces the brain volume that would be affected but makes the scalp a shorter circuit. Moreover the positioning of the electrodes so that the current flows parallel to the central sulcus might have exasperate variability between participant’s response to the modulation, because it has been suggested to produce non-uniform current directions across the cortical surface(29). Thus, the arguably neglectable effect size we found does not challenge tDCS in general, but our specific montage in particular. We are nevertheless keen to share our experience with our montage because we feel it has been an instructive and sobering example of how risky it is to take small effects in tDCS at face value without adding some more participants to see if the effect consolidates.

Our results highlight the need for robust replication of positive tDCS results to better understand the efficacy of, and mechanisms involved in, non-invasive brain stimulation. Further studies should be carried out to explore the potential of focality of HD-tDCS protocols, and the use of Bayesian statistics can provide us with the means to provide evidence in favour of either an effect or HD-tDCS or a lack thereof.

## ACKNOWLEDGEMENT

We thank Tatjana Maskaljunas and Balint K. Lammes for helping with the data collection.

This research was supported by the Netherlands Organization for Scientific Research (056-13-017, NIHC to C.K., VIDI 452-14-015 to V. G.), and the European Research Council of the European Commission (ERC-StG-312511 to C.K.).

